# Differential Correlation of Serum BDNF and microRNA Content in Rats with Rapid or Late Onset of Heavy Alcohol Use

**DOI:** 10.1101/2019.12.18.881680

**Authors:** Yann Ehinger, Khanhky Phamluong, David Darevesky, Melanie Welman, Jeffrey J. Moffat, Samuel A. Sakhai, Ellanor L. Whiteley, Anthony L. Berger, Sophie Laguesse, Mehdi Farokhnia, Lorenzo Leggio, Marie Lordkipanidzé, Dorit Ron

## Abstract

Heavy alcohol use reduces the levels of the brain-derived neurotrophic factor (BDNF) in the prefrontal cortex of rodents through the upregulation of microRNAs targeting *BDNF* mRNA. In humans, an inverse correlation exists between circulating blood levels of BDNF and the severity of psychiatric disorders including alcohol abuse. Here, we set out to determine whether a history of heavy alcohol use produces comparable alterations in the blood of rats. We used an intermittent access to 20% alcohol using the 2-bottle choice paradigm (IA20%2BC), and measured circulating levels of BDNF protein and microRNAs in the serum of Long-Evans rats before and after 8-weeks of excessive alcohol intake. We observed that the drinking profile of heavy alcohol users is not unified; Whereas 70% of the rats gradually escalate their alcohol intake (Late Onset), 30% of alcohol users exhibit a very Rapid Onset of drinking (Rapid Onset). We found that serum BDNF levels are negatively correlated with alcohol intake in both Rapid Onset and Late Onset rats. In contrast, increased expression of the microRNAs (miRs) targeting BDNF, miR30a-5p, miR-195-5p, miR191-5p and miR206-3p, was detected only in the Rapid Onset rats. Finally, we report that the alcohol-dependent molecular changes are not due to alterations in platelet number. Our data suggest that rats exhibit both Late and Rapid Onset of alcohol intake. We further show that heavy alcohol use produces comparable changes in BDNF protein levels in both groups. However, circulating microRNAs are responsive to alcohol only in the Rapid Onset rats.

## Introduction

Alcohol use disorder (AUD) is a detrimental health concern that affects 15% of the population in the US (Grant et al., 2015; WHO, 2014). Although the majority of people consume alcohol throughout their lifetime, only a small subset of them develop AUD. This conundrum suggests the existence of endogenous factors that protect the majority of the population from developing alcohol addiction.

One such endogenous protective factor is the brain derived neurotrophic factor (BDNF). BDNF belongs to the nerve growth factor (NGF) family of neurotrophic factors (Bothwell, 2014) which signals through tropomyosin-related kinase B (TrkB) receptor (Minichiello, 2009). BDNF binding to TrkB results in the activation of the mitogen activated protein kinase (MAPK)/extracellular signal regulated kinases 1 and 2 (ERK1/2), phospholipase C gamma (PLC gamma), and phosphoinositol 3-kinase (PI3K) pathways, and to the initiation of transcriptional and translational machineries (Huang and Reichardt, 2003; Leal et al., 2014; Ruiz et al., 2014). The BDNF/TrkB pathway plays an important role in central nervous system (CNS) development including neuronal differentiation and synapse formation (Park and Poo, 2013). In the adult brain, BDNF/TrkB signaling participates in synaptic and structural plasticity (Minichiello, 2009; Panja and Bramham, 2014), as well as learning and memory (Bekinschtein et al., 2014). In contrast, binding of BDNF or its immature proBDNF form to the low affinity p75 Neurotrophin receptor (p75NTR) activates a different set of targets including c-Jun amino terminal kinase (JNK), RhoA and nuclear factor kappa-light-chain-enhancer of activated B cells (NFkB) (Kraemer et al., 2014b), that in turn reduce spine complexity and density (Zagrebelsky et al., 2005), induce apoptosis (Kraemer et al., 2014a; Teng et al., 2005), and facilitate long term depression (Woo et al., 2005).

Over more than a decade, we and others observed that the BDNF system interacts with alcohol in a unique way (Logrip et al., 2015; Ron and Berger, 2018). We found that moderate but not excessive intake of alcohol increases the expression of BDNF in the dorsal striatum and more specifically in the dorsolateral striatum (DLS) of rodents (Jeanblanc et al., 2009; Logrip et al., 2009; McGough et al., 2004) resulting in the activation of TrkB/ERK1/2 signaling (Jeanblanc et al., 2013; Logrip et al., 2009; McGough et al., 2004), and in the induction of the dopamine D3 receptor (Jeanblanc et al., 2006) and preprodynorphin (Logrip et al., 2008) gene expression, which in turn keeps alcohol drinking in moderation. We further showed that dysregulation of the normal corticostriatal BDNF signaling drives the development and maintenance of rodents’ heavy alcohol use (Darcq et al., 2016; Darcq et al., 2015; Logrip et al., 2009; Warnault et al., 2016). For instance, we found that the development of excessive alcohol drinking is associated with increased contribution of p75NTR signaling in the DLS (Darcq et al., 2016), and in lowered *BDNF* levels in the prefrontal cortex (PFC) of rodents (Darcq et al., 2015; Logrip et al., 2009). In line with these findings, Tapocik et al. found that BDNF levels were reduced in rats that were made physically dependent on alcohol (Tapocik et al., 2014). We, and Tapocik et al., further showed that the alcohol-dependent suppression of BDNF expression in the mPFC is mediated through the induction of miR30-5p (Darcq et al., 2015), and miR206 (Tapocik et al., 2014) expression. In addition, alcohol-preferring rats display lower innate levels of BDNF in the bed nucleus of the stria terminialis (BNST) as well as in amygdalar regions as compared with alcohol non-preferring rats (Prakash et al., 2008; Raivio et al., 2014). Dysregulation of BDNF signaling has also been associated with alcohol withdrawal-induced anxiety-like behaviors in rats (Pandey et al., 2006). Finally, we found that a single point mutation in the BDNF gene which produces a valine to methionine substitution results in compulsive heavy alcohol use despite negative consequences in mice (Warnault et al., 2016). Interestingly, this single point mutation is also associated with an early onset of relapse in alcoholics undergoing treatment (Wojnar et al., 2009), and with an increased likelihood to respond to alcohol cues in adolescents (Nees et al., 2015). The Val66Met BDNF polymorphism can also predict the level of alcohol intake in alcohol-dependent patients when assessed together with polymorphisms for dopaminergic genes (Klimkiewicz et al., 2017). Together, these data suggest that BDNF is part of a homeostatic pathway that keeps alcohol intake in moderation, and that breakdown of this protective mechanism drives development and maintenance of AUD.

Similar to AUD, disruption of normal BDNF function has been associated with numerous of psychiatric disorders including depression, anxiety, stress and schizophrenia (Castren, 2014). Interestingly, a number of reports suggest that circuiting BDNF levels are inversely correlated with the severity of psychiatric disorders (Cattaneo et al., 2016). Similarly, humans with AUD have reduced serum BDNF content compared to healthy controls (Heberlein et al., 2010; Nubukpo et al., 2017; Silva-Pena et al., 2019; Zanardini et al., 2011; Zhou et al., 2018), which are still detected in abstinent alcoholics (Garcia-Marchena et al., 2017). Lower BDNF content has also been correlated with greater reported alcohol withdrawal severity in alcoholics (Costa et al., 2011; Heberlein et al., 2010). Finally, higher levels of BDNF was detected in alcoholics following 6 months of abstinence (Costa et al., 2011) suggesting that BDNF levels have been restored.

As disruption of normal BDNF signaling in the brain promotes the development of heavy alcohol use in rodents and possibly in humans (Ron and Berger, 2018), and as lowered serum BDNF levels are associated with AUD, we set out to determine if the same is true for the BDNF machinery in the peripheral blood serum of rats. We further determined whether the decrease in BDNF serum content is associated with alterations in the levels of microRNAs targeting BDNF.

## Methods

### Reagents

BDNF E_max_®ImmunoAssay System kit (cat. #G7610) was purchased from Promega (Madison, WI). BCA™ Protein Assay (cat. #23227) was purchased from Thermo Fisher Scientific (Waltham, MA). miRNeasy Serum/Plasma Kit (cat. #217184), miRCURY LNA RT kit (cat. #339340), and miRCury LNA SYBR Green master mix (cat. #339346) were purchased from Qiagen (Germantown, MD). Ethyl alcohol (190 proof) was purchased from VWR (Radnor, PA).

### Animals

Male adult Long-Evans rats (∼8-10 weeks of age) weighing approximately 250-300 grams were obtained from Harlan Labs. Animals were individually housed on paper-chip bedding (Teklad #7084), under a 12 hour light-dark cycle (lights on 0600 to 1800 h). Temperature and humidity were kept constant at 22 ± 2°C, and relative humidity was maintained at 50 ± 5%. Rats were allowed access to food (Teklad Global Diet #2918) and tap water *ad libitum*. All animal procedures were approved by the University of California, San Francisco Institutional Animal Care and Use Committee and were conducted in agreement with the Association for Assessment and Accreditation of Laboratory Animal Care (AAALAC, UCSF). Two independent cohorts were used for serum BDNF content and microRNA analysis. A third cohort of rats was used for platelet number analysis.

### Preparation of alcohol solution

Alcohol solution was prepared from ethyl alcohol absolute anhydrous (190 proof) diluted 20% (v/v) in tap water.

### Alcohol drinking paradigm

The intermittent-access to 20% alcohol two-bottle choice drinking procedure (IA20%-2BC) was conducted as previously described (Carnicella et al., 2009). Specifically, rats were given access to one bottle of 20% alcohol (v/v) in tap water and one bottle of water for 24 hours. Control mice had access to water only. Drinking sessions started at 12:00PM on Monday, Wednesday and Friday, with 24- or 48 hours (weekend) of alcohol-deprivation periods in which rats consumed only water. The placement (left or right) of water or alcohol solution was alternated between each session to control for side preference. Water and alcohol bottles were weighed at the beginning and end of each alcohol drinking session. Rats were weighed once a week.

### Serum blood collection

Rats were bled via the lateral tail vein before the initiation of the IA20%-2BC protocol and 24 hours after the final IA20%-2BC session (Fig. 1A). Animals were consistently bled in the morning. Whole blood was kept for one hour at room temperature followed by centrifugation at 2000 x g for 10 minutes at 4°C. Room temperature incubation time was strictly monitored, as clotting can influence serum levels of BDNF (Maffioletti et al., 2014). The resulting supernatant was aliquoted, snap frozen to minimize RNA degradation, and stored at -80°C for later ELISA and microRNAs expression analysis.

**Figure 1.**
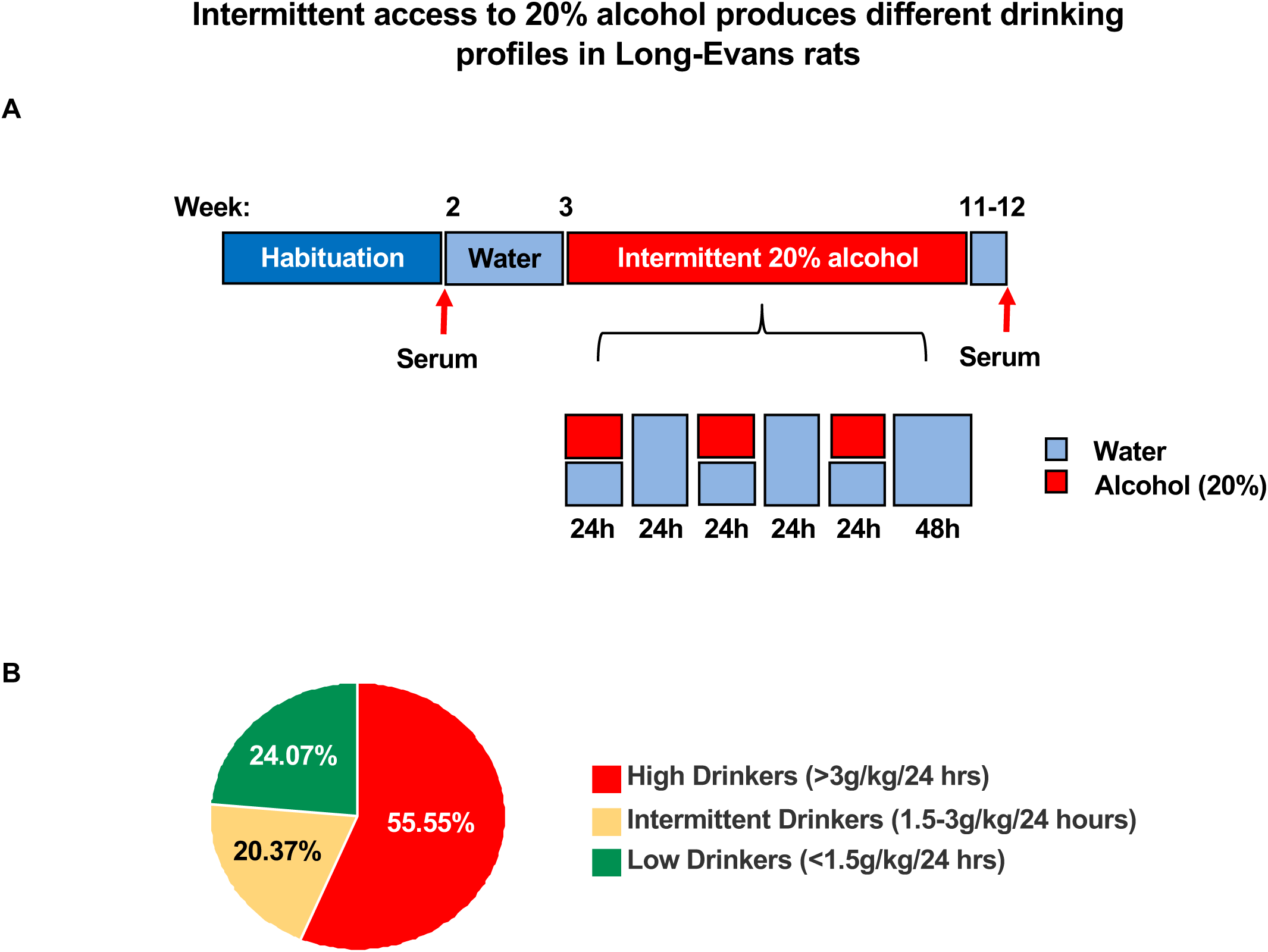
Intermittent access to 20% alcohol produces different drinking profiles in Long-Evans rats. **(A)** Timeline of the alcohol drinking paradigm and serum collection days. (**B**) Pie chart depicts the percentage of low drinkers which consumed an average of 0.97 ± 0.065 g/kg/24 hour, intermediate drinkers which drank an average of 2.34 ± 0.085 g/kg/24 hour and high drinkers which drank an average of 4.27 ± 0.116 g/kg/24 hour. n = 13 low drinkers, 11 intermediate drinkers, n=30 high drinkers.

### Enzyme-linked immunosorbent assay (ELISA)

ELISA assay was conducted according to manufacturer’s instructions with minor modifications. Specifically, 96-well plates were coated with 100 μl of monoclonal anti-BDNF antibody solution and incubated covered overnight at 4°C without shaking. Wells were washed once with TBST (20 mM Tris–HCl (pH 7.6), 150 mM NaCl, 0.05% Tween 20), then incubated with 200 μl of a Block & Sample buffer 1X for 1 hour at room temperature without shaking. Wells were washed once with TBST. Standard curves were prepared by diluting BDNF standards in Block & Sample buffer, with six serial 1:2 dilutions. A blank control was added for each plate. Samples were diluted in the same buffer (1:40) and added in triplicate. Plates were incubated for 4 hours at room temperature with shaking, and subsequently washed five times with TBST. One hundred μl of anti-BDNF polyclonal antibody solution was added and plates were incubated overnight at 4°C with shaking. After five washes with TBST, 100 μl of secondary HRP conjugate antibody solution was added and plates were incubated for 1 hour at room temperature with shaking. Wells were washed five times with TBST. Subsequently, 100 μl of 3,3’,5,5’-tetramethylbenzidine (TMB) One Solution was added to each well, and plates were incubated for 30 minutes at room temperature. Reactions were stopped with 100 μl of 1M HCl, and absorbance at 450 nm was measured with an automated microplate reader within 30 minutes. Standard curves were plotted for each plate. Triplicates were averaged and values were corrected for total amount of protein in the sample as determined by BCA protein assay.

### MicroRNA extraction and cDNA synthesis

MicroRNAs were extracted from the blood serum fraction using the miRNeasy Serum/Plasma Kit according to the manufacturer’s instructions. RNA yield and purity were evaluated using a nanodrop ND-1000 spectrophotometer (ThermoFisher Scientific, Waltham, MA). cDNA synthesis was performed using the miRCURY LNA RT kit according to the manufacturer’s instructions, starting with 150 ng of RNA and 5X reaction mix. Enzyme and nuclease-free water was added to a final volume of 20 μl. RNA spike-in of Unisp6 was added to each sample to monitor the efficiency of the reverse transcription reaction.

### Real-time quantitative reverse transcriptase PCR (qRT-PCR)

qRT-PCR was performed using miRCURY LNA SYBR Green master mix according to the manufacturer’s instructions, and experiments were run on an AB7900HT fast real-time PCR instrument (Applied Biosystems, Foster City, CA). Each data point represents an average of three replicates. Relative microRNA expression for each sample was determined by the following calculation; First, the average of cycle threshold (CT) of the miRNA of interest was subtracted from the CT of the control 5S rRNA to generate ΔCT. ΔCTs were then subtracted from the average of control ΔCT values for each sample to generate the ΔΔCT. Fold changes were then calculated as 2^-ΔΔCT^+/-S.E.M.

The following locked nucleic acid (LNA) PCR primers were used:

miR30a-5p: 5’-UGUAAACAUCCUCGACUGGAAG-3’, (Qiagen, cat. #YP00205695);

miR195-5p: 5’-UAGCAGCACAGAAAUAUUGGC-3’, (Qiagen, cat. #YP00205869);

miR206-3p: 5’-UGGAAUGUAAGGAAGUGUGUGG-3’, (Qiagen, cat. #YP00206073);

miR191-5p: 5’-CAACGGAAUCCCAAAAGCAGCUG-3’, (Qiagen, cat. #YP00204306);

miR34c-5p: 5’-AGGCAGUGUAGUUAGCUGAUUGC-3’, (Qiagen, cat. #YP00205659);

Controls: 5S rRNA (Qiagen, cat. #YP00203906) and UniSp6 (Qiagen, cat. #YP00203954).

### Platelet number analysis

Whole blood was collected from the tail artery directly into EDTA blood collection tubes (K2 EDTA 10.8 mg, BD Vacutainer) using a 25G needle (BD Vacutainer). An aliquot of 500 μl was used for complete blood count analysis. Platelet counts were determined using a Beckman Coulter Hematology Analyzer Ac·T 5diff AL (Beckman Coulter, Orange County, CA, USA). Experimenters conducting the analysis were blind to the group assignment.

### Statistical analyses

Statistical analyses were performed using SigmaStat 4.0 statistical software (San Jose, CA.). Statistical significance was determined by using two-way ANOVA with repeated measures, Post-hoc Student-Newman-Keus (SNK) test, and two-tailed t-test as indicated in the Figure Legends. K-means clustering analysis was used as described in (Krahe et al., 2011) to determine the most parsimonious number of clusters for partitioning alcohol drinkers into behavior phenotypes based on the average alcohol intake (in g/kg) of the first four and last four drinking sessions. This was performed in an unsupervised manner by running K-means clustering on the data with different values of *k* ranging from 1 to 10, inclusive. The sum-of-squares error (SSE) for each value of *k* was calculated and the elbow criterion (James et al., 2013) was used to identify the corresponding *k* at which the asymptote plateaued (See Suppl. Fig. 1 for plot of number of clusters (*k*) vs SSE). K-means was then performed on the drinking data using the value of *k* determined with the aforementioned method. All clustering analyses were performed with Python v3.7 using the SciPy stack. Significance was set at p < 0.05.

## Results

### Animals that consume high levels of alcohol exhibited Late or Rapid Onset of heavy alcohol use

To investigate the association between serum BDNF and heavy alcohol use, we used Long-Evans rats as a model system and measured BDNF protein in blood serum prior to and at the end of an 8 week IA20%-2BC paradigm, a procedure that models episodic heavy alcohol drinking in humans (Carnicella et al., 2014) (Figure 1A).

We observed that out of the 54 rats used in the study, only 55.55 % of animals (n=30) were heavy drinkers consuming more than 3 g/kg/24 hours with an average of 4.27 +/-0.16 g/kg/24 hours at the last 4 sessions, which corresponds to blood alcohol concentration (BAC) of above 80 mg/dl or higher (Carnicella et al., 2014) (Figure 1B). This group of rats were used for the study. Rats that consumed less than 1.5 g/kg/24 hours and an average of 0.97 +/-0.065 g/kg/24 hours were considered low drinkers and accounted for 24.07% of the animals (n=13). The remaining 20.37% of animals (n=11) were intermediate drinkers consuming alcohol more than 1.5 g/kg/24 hours but less than 3 g/kg/24 hours with an average of 2.34 +/-0.085 g/kg/24 hours (Figure 1B).

Surprisingly, heavy alcohol users could be segregated into two distinct group; 70% of the animals (n=21) gradually escalated their alcohol use, drinking an average of 1.06 +/-0.152 g/kg/24 hours on week 1 vs. 4.23 +/-0.318 g/kg/24 hours on week 8 (Figure 2A-C, red). These rats were defined as Late Onset animals. In contrast, 30% of the animals (n=9) consumed high levels of alcohol, even on the first session of drinking (Figure 2A-B,D, blue), and the amount of alcohol consumed throughout the 8 weeks drinking period remained constant (4.92 +/-0.304 g/kg/24 hours on week 1 vs. 4.45 +/-0.445 g/kg/24 hours on week 8) (Figure 2B,D, blue). These rats were defined as Rapid Onset animals.

**Figure 2.**
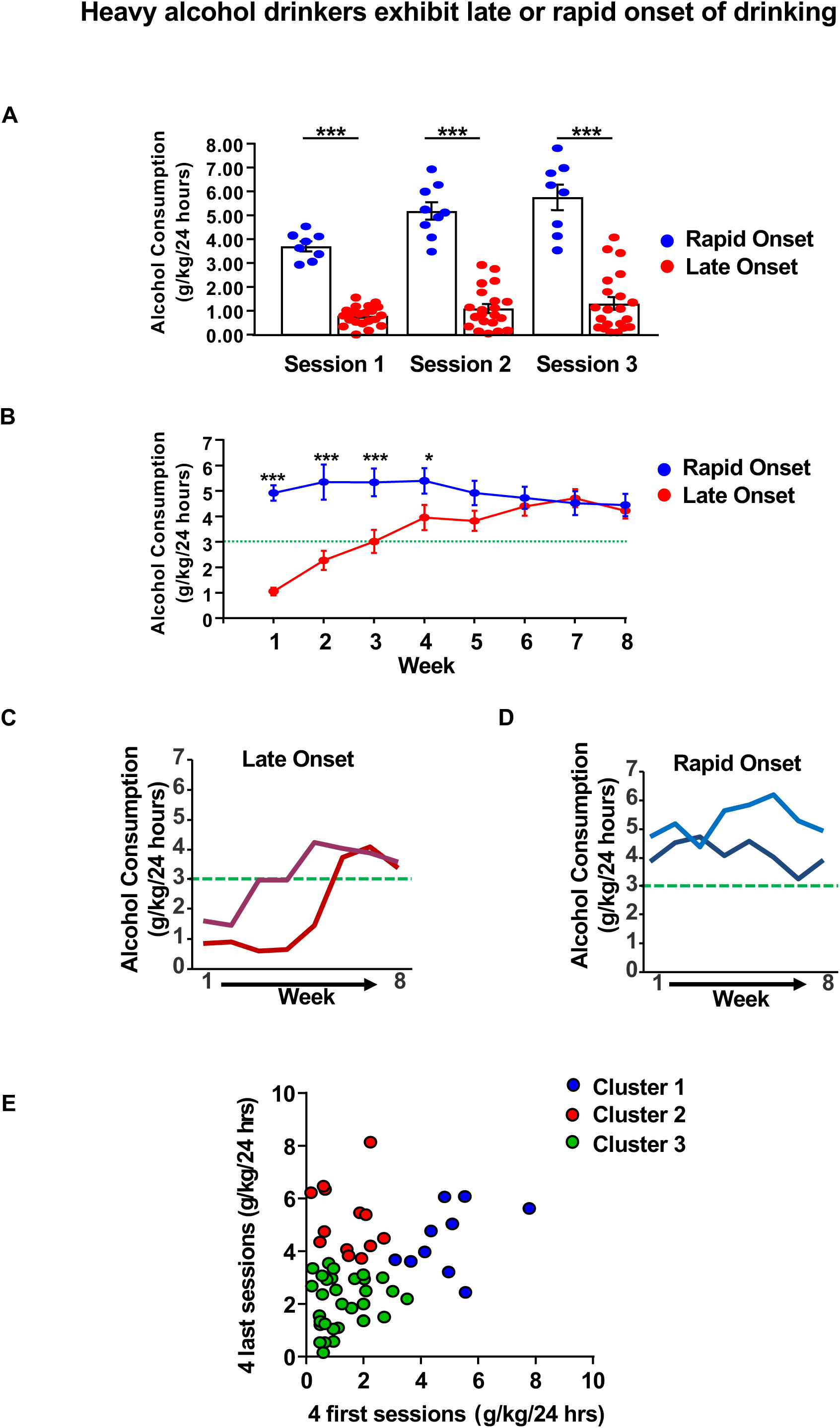
Heavy alcohol drinkers exhibit a Late or Rapid Onset of drinking. (**A-**B) Graphs depict the average alcohol consumption +/- SEM of **Late Onset** (red) and **Rapid Onset** (blue) rats during the first 3 sessions (**A**), and during the 8 weeks of drinking period, i.e. 24-26 drinking sessions (**B**). (**C-D**) Representative drinking profiles of Late Onset (C, red) and Rapid Onset (**D**, blue) rats. **(E)** Drinking data clustered by K-means clustering (k=3) with k determined by the “elbow method” (see methods for details). Animals starting and ending at a low alcohol intake level in the first four sessions and last four sessions, clustered into one group (cluster 3, green). High drinkers are divided into two distinct clusters: one group starts at a low intake level in the first four sessions (cluster 2, red) and second one starting at a high intake level in the first four sessions (cluster 1, blue). (**A)** Rapid onset n=9, Late Onset n=21, two-way repeated measures ANOVA (one factor repeated) with Student-Newman Keuls post-hoc test: Late Onset vs. Rapid Onset p < 0.001, in sessions 1-3. **(B)** Rapid Onset n=9, **Late Onset** n=21, two-way repeated measures ANOVA (one factor repeated) with Student-Newman-Keuls post-hoc test: week 1 (**Late Onset vs. Rapid Onset**): p < 0.001, week 2-3: p < 0.001, week 4: p < 0.05, week 5 to 8. (**C-D**) n=2 per group. (**E**) n=30. * p< 0.05, *** p< 0.001.

In order to determine the number of clusters with an unsupervised approach, K-means clustering analysis (Krahe et al., 2011) was performed on the drinking data (the first four and last four drinking sessions). This approach confirmed the segregation of high drinkers in two groups (Figure 2E); One cluster of animals exhibited Rapid Onset of drinking (Figure 2E, cluster 1, blue), and another cluster corresponded to Late Onset of drinking (Figure 2E, cluster 2, red). Animals drinking on average below ∼3.5g/kg over the first four and last four drinking sessions were clustered into a third group (Figure 2E, cluster 3, green).

Together, these data indicate that rats exhibit two markedly different profiles of excessive drinking; rats whose alcohol intake gradually escalate, and animals that consume large quantities of alcohol at the initiation of the drinking paradigm.

### Late onset and Rapid Onset heavy alcohol drinkers exhibit a decrease in serum BDNF

To measure BDNF content in the blood serum before and after a period of heavy alcohol use, we compared the basal levels of serum BDNF protein to the levels of the neurotrophic factor after 8 weeks (24-26 sessions) of IA20%-2BC (Figure 1A). We found that similar to the mPFC (Darcq et al., 2015; Tapocik et al., 2014), heavy alcohol use reduces serum BDNF levels (Figure 3A). We then examined whether Late Onset and Rapid Onset rats exhibit a similar profile of BDNF quantity before and after excessive alcohol consumption. As shown in Figure 3B, the reduction in BDNF protein content was observed in both Rapid Onset and Late Onset rats. We next assessed whether the levels of serum BDNF are associated with other factors irrespective of the drinking phenotypes. To do so, we correlated serum BDNF levels with water consumption and found that 8 weeks of water intake had no effect on BDNF levels (Figure 3C). Together, these results show that a history of heavy alcohol use produces a reduction in serum BDNF content. Our data further suggests that low BDNF protein levels are detected after a history of heavy alcohol intake regardless of the rats’ drinking profile.

**Figure 3.**
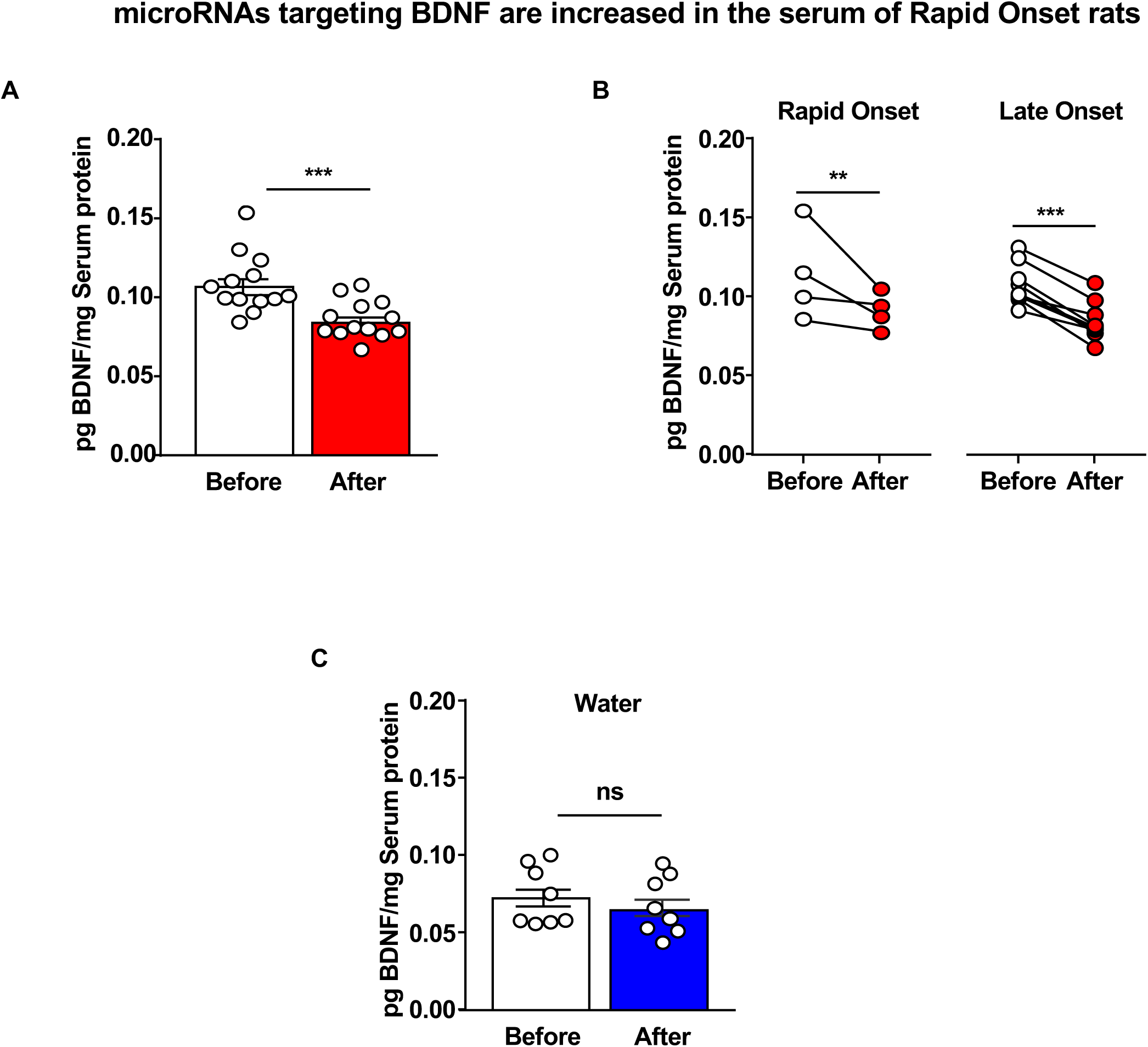
Serum BDNF protein levels are reduced by alcohol in Late Onset and Rapid Onset rats. Rats underwent an IA20%-2BC drinking paradigm for 8 weeks. Animals were bled via lateral tail vein one week before the initiation of the drinking paradigm (basal) and 24 hours after the final drinking session. Timeline is depicted in Figure 1. BDNF protein level was measured by ELISA and normalized to the mean of total blood serum protein. (**A**) Bar graph depicts individual data point and mean +/- SEM of BDNF protein content before (white) and at the end of the last 24 hours of withdrawal (red). (**B**) Individual data points of serum BDNF content from Late Onset (*right*) and Rapid Onset (*left*) animals. (**C**) Bar graph depicts average data points and mean +/- SEM of serum BDNF content before (white) and after (blue) 8 weeks of water intake. (**A**) n=13, two-tailed paired t-test: t(8) = 9.15, p < 0.001. (**B**) n=4 Rapid Onset, n=9 Late Onset, two-way repeated measures ANOVA (one repeated factor) with Student-Newman-Keuls post-hoc test: Rapid Onset before vs. after p < 0.01, Late Onset before vs. after p < 0.001. (**C**) n=8, two-tailed paired t-test: t(7) = 1.17, p = 0.28. ** p < 0.01. *** p < 0.001. ns: none-significant.

### microRNAs targeting BDNF are upregulated in the serum of Rapid Onset alcohol users

MicroRNAs have the capacity of targeting hundreds of mRNAs for degradation and/or translational repression (Qureshi and Mehler, 2012). In addition, several microRNAs can target the same gene (Qureshi and Mehler, 2012). Strikingly, BDNF is a target of numerous microRNAs including miR30a-5p (Darcq et al., 2015; Mellios et al., 2009; Mellios et al., 2008), miR195 (Mellios et al., 2009; Mellios et al., 2008), miR124 (Chandrasekar and Dreyer, 2009), and miR206 (Tapocik et al., 2014).

As BDNF expression is tightly regulated by numerous microRNAs, and as microRNAs are found in serum (Cortez and Calin, 2009), we measured the levels of miR30a-5p, miR195-5p, miR206-3p, and miR191-5p before and after IA20%2BC in Rapid Onset and Late Onset rats. We found that the basal level of the microRNAs is low in the serum of animals prior to alcohol exposure. As shown in Figure 4A,C,E,G, IA20%2BC produced a significant increase in the level of all 4 tested microRNAs ranging from 2 fold increase (miR191-5p) to 20-fold increase (miR206-3p). Strikingly however, all 4 microRNAs were increased by alcohol only in the Rapid Onset animals but not in Late Onset animals (Figure 4B,D,F,H). We then measured the levels of miR134c-5p which is found in serum (Jin et al., 2018) but has not been associated with the regulation of BDNF expression. As shown in Figure 5, there was no correlation between miR134-5p levels and alcohol intake. Together, these data suggest that microRNAs miR30a-5p, miR195-5p, miR206-3p, and miR191-5p known to target BDNF in the brain are robustly increased by alcohol in serum of Rapid Onset animals.

**Figure 4.**
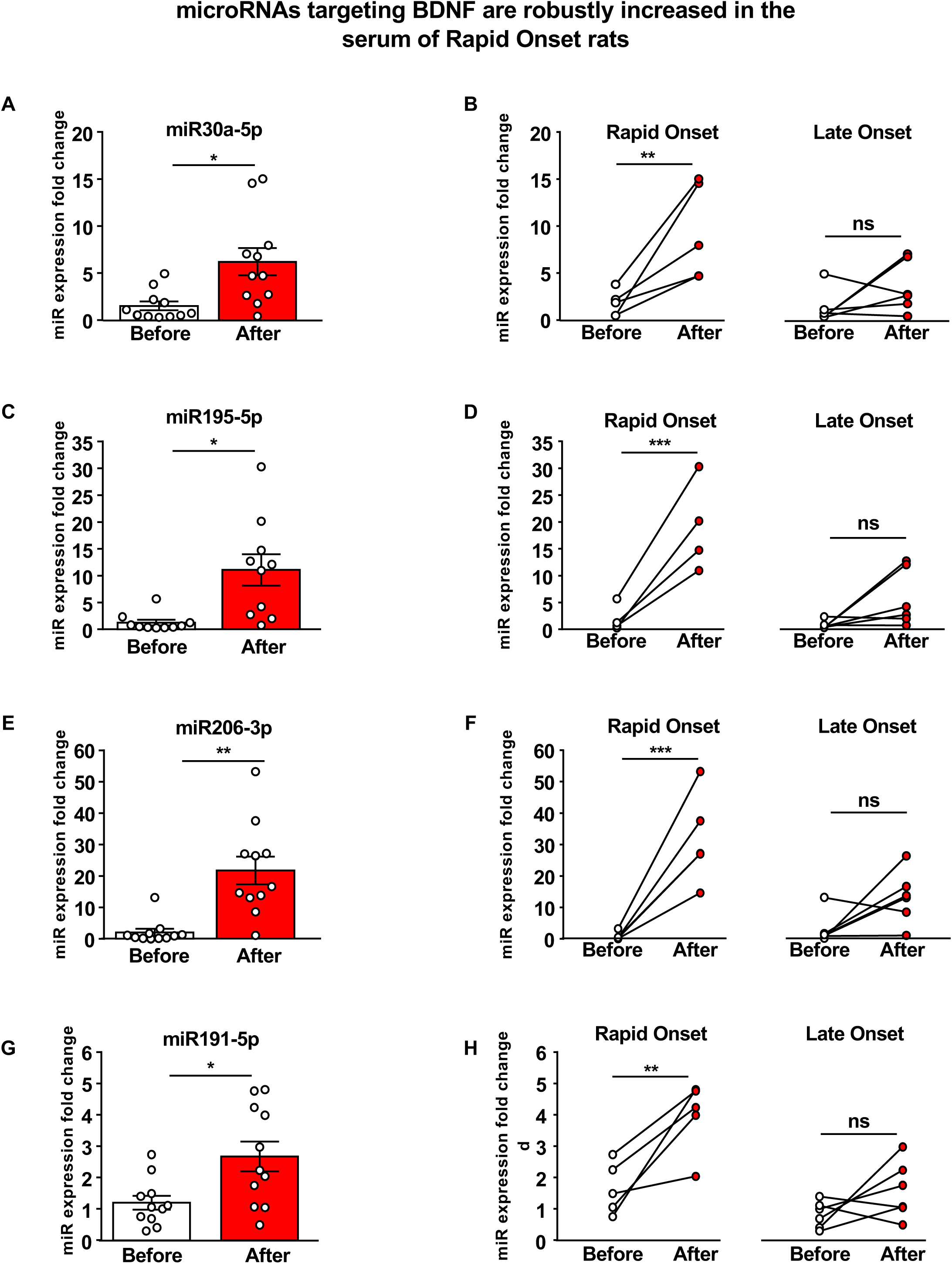
microRNAs targeting BDNF are increased in the serum of Rapid Onset rats. Rats underwent an IA20%-2BC drinking paradigm for 8 weeks. Animals were bled via lateral tail vein one week before the initiation of the drinking paradigm (basal) and 24 hours after the final drinking session. Timeline is depicted in Figure 1. The levels of microRNAs miR30a-5p (**A-B**), miR195-5p (**C-D**), miR206-3p (**E-F**) and miR191-5p (**G-H**) were measured by qRT-PCR. (**A**,**C**,**E**,**G**) Bar graphs depict individual data points and mean +/- SEM of microRNAs before (white) and after (red) alcohol use. (**B**,**D**,**F**,**H**) Individual data points of serum microRNA levels from Late Onset (*right*) and Rapid Onset (*left*) animals. (**A**) n=11, two-tailed paired t-test: t(5) = 3.55, p < 0.05. (**B**) n=5 Rapid Onset, n=6 Late Onset, two-way repeated measures ANOVA (one factor repeated) with Student-Newman-Keuls post-hoc test: Rapid Onset before vs. after: p <0.01, Late Onset before vs. after: p=0.22. (**C**) n=10, two-tailed paired t-test: t(4) = 5.32, p < 0.05. (**D**) n=4 Rapid Onset, n=6 Late Onset, two-way repeated measures ANOVA (one factor repeated) with Student-Newman-Keuls post-hoc test: Rapid Onset before vs. after p < 0.001, Late Onset before vs. after p = 0.08. (**E**) n=11, two-tailed paired t-test: t(5) = 5.22, p < 0.01. (**F**) n=5 Rapid Onset, n=6 Late Onset, two-way repeated measures ANOVA (one factor repeated) with Student-Newman-Keuls post-hoc test: Rapid Onset before vs. after p < 0.001, Late Onset before vs. after p=0.07. (G) n=11, two-tailed paired t-test: t(5) = 4.00, p < 0.05. (**H**) n=5 Rapid Onset, n=6, two-way repeated measures ANOVA (one factor repeated) with Student-Newman-Keuls post-hoc test: Rapid Onset before vs. after p < 0.01, Late Onset before vs. after p = 0.16. * p< 0.05, ** p< 0.01, *** p< 0.001. ns: none-significance.

**Figure 5.**
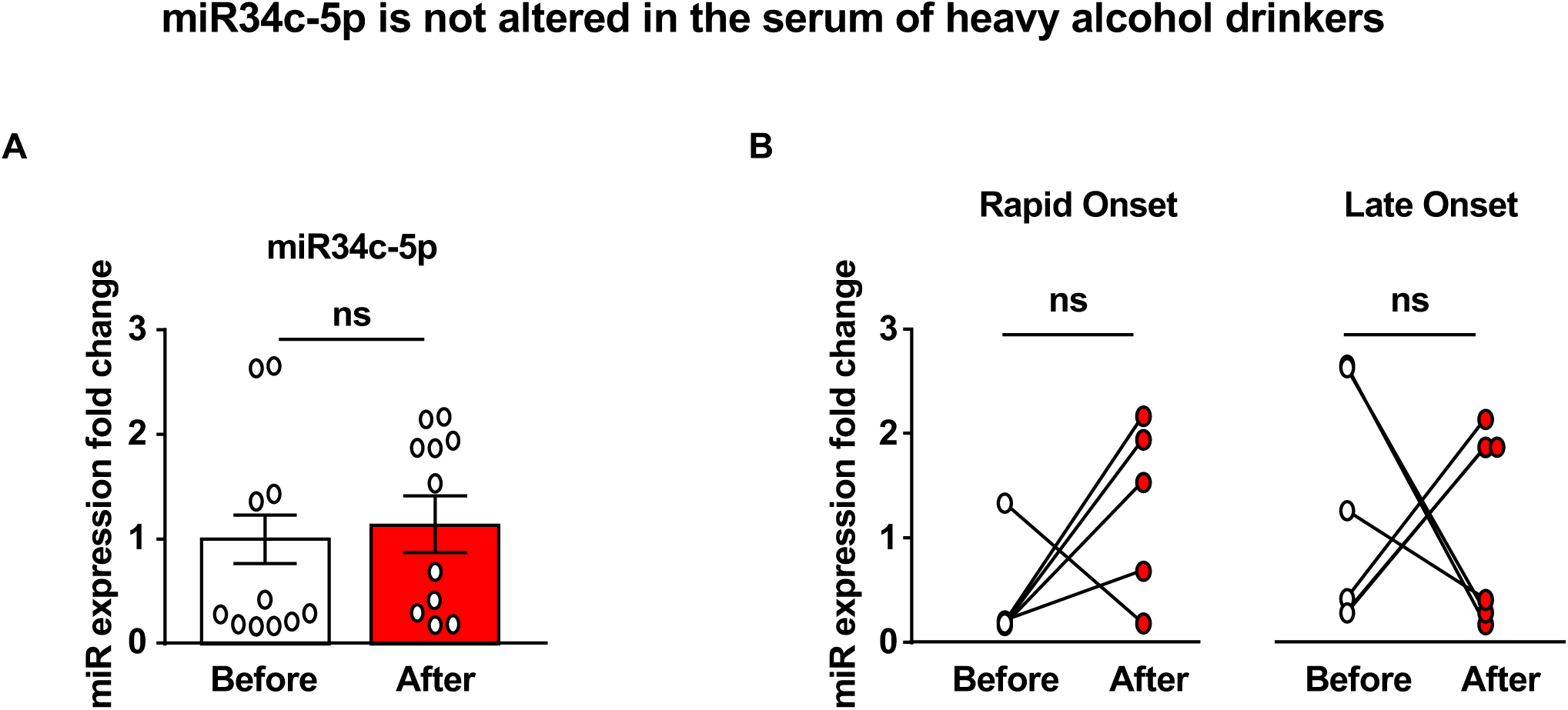
miR34c-5p is not altered in the serum of heavy alcohol drinkers. Experimental design is depicted in Figure 1A, and described in Figure 4. Animals were bled via lateral tail vein one week before the initiation of the drinking paradigm (basal) and 24 hours after the final drinking session. miR34c-5p was measured by qRT-PCR. (**A**) Bar graph depicts individual data point and mean +/- SEM of miR34c-5p before (white) and after (red) alcohol use. **(B)** Individual data points from Late Onset (*right*) and Rapid Onset (*left*) animals. (**A**) n=11, two-tailed paired t-test: t(11) = 0.64, p = 0.54. (B) n=6 Late Onset n=5, Rapid Onset, two-way repeated measures ANOVA with Bonferroni post-hoc test: Rapid Onset before vs. after p=0.27, Late Onset before vs. after p=0.83. ns: non-significant.

### Platelet number is not altered by heavy alcohol use

Platelets represent the largest pool of BDNF in blood (Chacon-Fernandez et al., 2016). Secretion of BDNF by activated platelets is thus an important regulator of its bioavailability in serum (Fujimura et al., 2002). Platelets are also a source of a large number of circulating microRNAs (Sunderland et al., 2017). In humans, chronic alcoholism can cause thrombocytopenia, i.e. low platelet count (Eichner, 1973). Thus, we determined whether the alterations in BDNF and miRs level in response to heavy alcohol use in rats could be due to a lower platelet number in the alcohol consuming rats. To do so, the platelet count was analyzed before and after 8 weeks of IA20%2BC. We found platelet counts to be unaffected by 8 weeks of heavy alcohol use, regardless of the rapidity of onset of heavy alcohol consumption (Figure 6).

**Figure 6.**
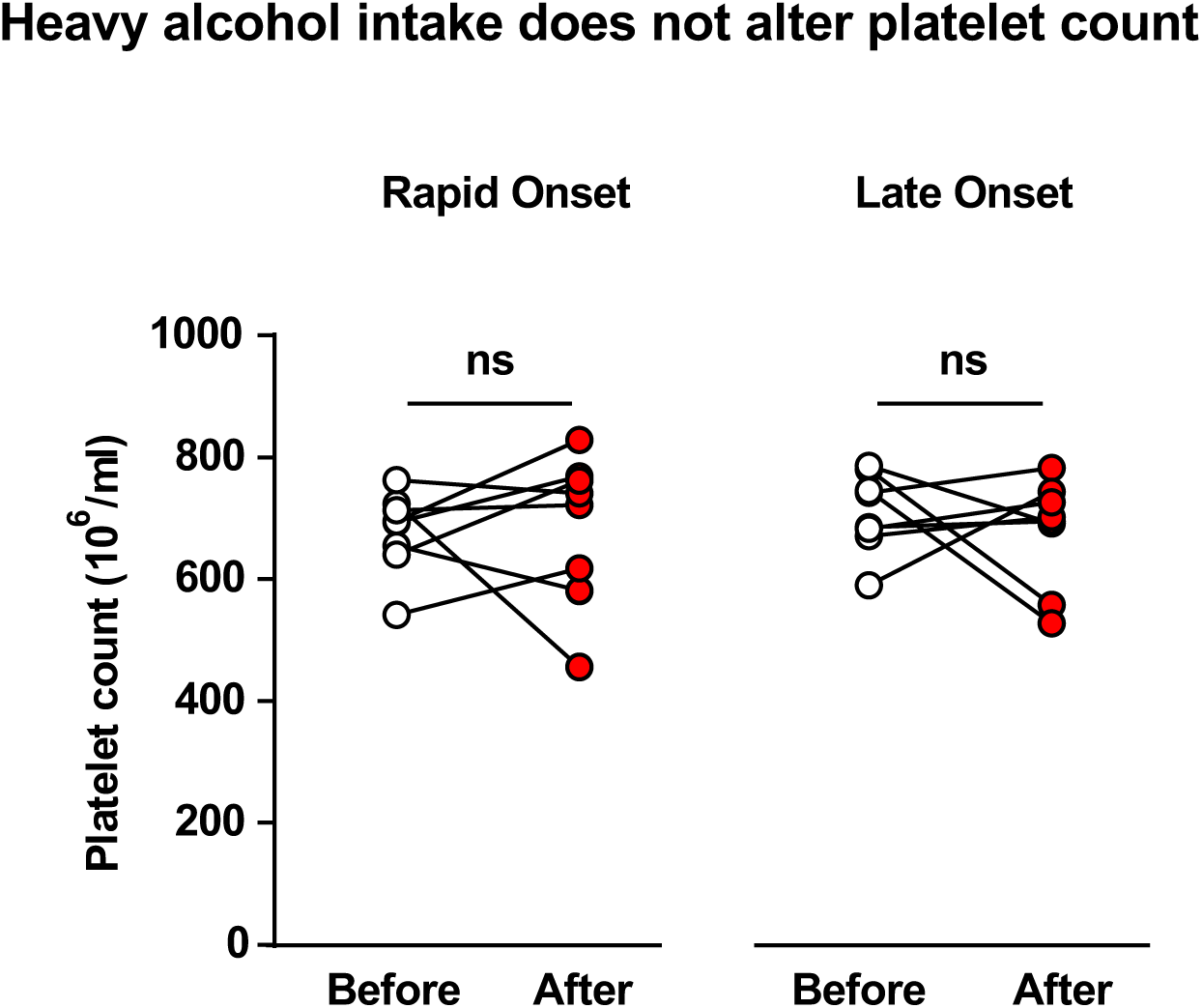
Heavy alcohol intake does not alter platelet count. Experimental design is depicted in Figure 1A. Animals were bled via tail artery one week before the initiation of the drinking paradigm (basal) and 24 hours after the final drinking session. No changes were observed in platelet count before and after 8 weeks of IA20%2BC. n=8 Late Onset, n=8 Rapid Onset. Two-way repeated measures ANOVA; effect of drinking profile: F_1,14_=0.1841, p=0.6744; effect of time F_1,14_=0.1459, p=0.7083; interaction: F_1,14_=0.3293, p=0.5752.

## Discussion

Our study consists of three major findings: We report that IA20%2BC produces different profiles of heavy alcohol drinking. Specifically, we found that whereas the majority of rats gradually escalate their alcohol intake, a portion the animals consume large quantities of alcohol even at the initiation of the alcohol drinking regimen. We further show that serum BDNF levels are reduced in heavy alcohol drinking rats regardless of their alcohol drinking profile. We also report that alcohol increases the level of microRNAs targeting BDNF only in the serum of rats that exhibit Rapid Onset of drinking. Finally, we show that the alterations in BDNF levels as well as the expression of microRNAs targeting BDNF in response to excessive alcohol use are not due to changes in the number of platelets.

### Rats exhibit Late Onset and Rapid Onset drinking profiles

AUD is characterized, in part, by excessive alcohol intake (Edwards et al., 2013; Enoch and Goldman, 2002), and the IA20%2BC in rodents has been used as a reliable paradigm to probe for mechanisms that underlie the human disease (Carnicella et al., 2014). Here, we report that this regimen does not produce homogenous levels of intake. We observed that only half of the rats consumed over 3g/kg/24 hours, an amount of alcohol that generates BAC of over 80mg% (Carnicella et al., 2014). Surprisingly, we found that rats that drank high levels of alcohol can be further divided into two distinct groups: rats that start drinking low amounts of alcohol and escalate their alcohol drinking over a period of time (Late Onset), and rats with a very Rapid Onset of intake (Rapid Onset). This observation was detected in additional 4 cohorts of animals (data not shown), and was confirmed by an unsupervised K-means clustering analysis. It is plausible that the Late Onset and Rapid Onset animals have unique biological markups that underlie their drinking phenotypes. For example, it is possible that the genetic makeup of Long-Evans rats is not homogenous and that innate differences may be the root cause of the diverse drinking profiles. Furthermore, genetic or epigenetic makeup of the Rapid Onset heavy drinkers may resemble the alcohol preferring rats (P rats), whereas the low drinkers resemble the alcohol non-preferring rats (NP rats) (McBride et al., 2014). The Rapid Onset heavy alcohol users are of special interest as Rapid Onset heavy drinking animals may be an easily utilized rodent paradigm to model human individuals that show a Rapid Onset and sustained binge drinking of alcohol. In line with this possibility, Gowin et al. reported that binge drinking of alcohol is an early indicator of humans that are at risk of developing AUD (Gowin et al., 2017). Augier et al. showed that 15% of animals prefer alcohol over the highly rewarding substance, saccharine (Augier et al., 2018). The authors further showed that the vulnerable rats also exhibited addiction-like phenotypes such as consuming alcohol despite adverse consequences (Augier et al., 2018). Although this model has significant value, it is cumbersome and expensive to use. It is thus plausible that the Rapid Onset high drinking rats could be an easier model to study the addiction-prone population. Whether the Rapid Onset heavy alcohol users also exhibit other addiction-like phenotypes will be addressed in future studies. Regardless, differences in the drinking profiles of Long-Evans rats should be taken into consideration when investigating the neurobiological underpins of heavy alcohol use.

### BDNF protein levels are dysregulated in Late Onset and Rapid Onset rats

In this study, we show that serum BDNF becomes dysregulated after 8 weeks of IA20%-2BC. The decrease in BDNF levels were detected in the serum of heavy drinkers regardless of whether or not they rapidly or slowly escalate their alcohol use. These results mirror previous findings that dysregulation of BDNF physiology is associated with AUD phenotypes in rodents (Darcq et al., 2015; Logrip et al., 2015; Logrip et al., 2009; Tapocik et al., 2014; Warnault et al., 2016), as well as with the pathogenesis of AUD in humans (Ceballos and Sharma, 2016). The potential use of BDNF as a biomarker of AUD is worthy of more detailed exploration.

A caveat of our study is the fact that we measured the levels of BDNF at only two time points, before and after 8 weeks of drinking. A careful time course is required to determine whether the escalation in alcohol drinking in the Late Onset animals parallels a gradual reduction in serum BDNF content. Interestingly, a meta-analysis in patients with major depression demonstrated that BDNF levels in the blood are restored in patients that responded to an anti-depressant medication, whereas BDNF blood concentration remained unchanged in non-responders (Polyakova et al., 2015). These data suggest that BDNF in the blood is sensitive to a drug that targets the brain. Whether or not BDNF in blood cells has a biological function and whether alterations in serum BDNF levels by alcohol contribute to the disease are open questions. These questions are relevant not only to AUD, but also to other psychiatric disorders as serum BDNF is inversely correlated with depression (Molendijk et al., 2011), bipolar disorder (Fernandes et al., 2011), and schizophrenia (Rizos et al., 2008).

Megakaryocytes, the progenitor cells and are a major source of BDNF in platelets (Chacon-Fernandez et al., 2016). Specifically, megakaryocytes contain high levels of BDNF mRNA (Chacon-Fernandez et al., 2016). The BDNF protein is stored in platelets in α-granules (Chacon-Fernandez et al., 2016), and is secreted into the plasma upon platelet activation (Fujimura et al., 2002). Thus, it would be of interest to determine whether the BDNF production machinery in megakaryocytes and BDNF secretion mechanisms by platelets of Late Onset and Rapid Onset animals are altered by alcohol. BDNF is also expressed in T lymphocytes, B lymphocytes and monocytes (Linker et al., 2010; Linker et al., 2015), as well as in endothelial cells (Nakahashi et al., 2000)(Kermani and Hempstead, 2019). Future studies will determine whether alcohol alters BDNF’s levels in one or all of these cell-types.

### microRNAs targeting BDNF are upregulated only in Rapid Onset animals

We found that the levels of serum microRNAs targeting BDNF are relatively low in water consuming rats, however, miR30a-5p, miR195-5p, miR206-3p and miR191-5p expression were robustly upregulated by alcohol. In contrast, miR34c-5p, which does not target BDNF, was unaltered by alcohol. Strikingly, the increase in serum microRNAs was detected only in the Rapid Onset animals. This surprising result is yet another evidence that the Late Onset and Rapid Onset animals are genetically and/or epigenetically different.

Circulating microRNAs are transcribed in mononuclear blood cells, and packaged into exosomes which are released into the bloodstream and are internalized into recipient cells through an endocytosis process (Cortez and Calin, 2009). Serum microRNAs have been associated with numerous pathologic states (Cortez and Calin, 2009), including cancer (Mitchell et al., 2008), autoimmune diseases (Jin et al., 2018), as well as autism (Anitha and Thanseem, 2015). A small-scale study also suggested that microRNA levels are altered in the serums of human subjects suffering from AUD (Ignacio et al., 2015). It would be of interest to determine from which cell type the microRNAs are originating and through which mechanism alcohol produces a robust induction of their expression. It is likely that microRNAs target BDNF within the same cells however this possibility needs to be confirmed in future studies. Another remaining unanswered question is whether circulating microRNAs themselves have signaling properties, or are they merely a byproduct of cell homeostasis process.

Interestingly, the changes in miR30a-5p and miR206-3p in the serum of Rapid Onset rats is similar to the alterations detected in the mPFC of rodents (Darcq et al., 2015; Tapocik et al., 2014). Recently, Torres-Berrio et. al. reported that miR218 in the blood of mice is produced in the mPFC (Torres-Berrio et al., 2019). Thus, it is plausible that miRs targeting BDNF are produced in the mPFC transported by blood, where they can target BDNF mRNA in megakaryocytes, for example (Chacon-Fernandez et al., 2016). Thus, if the periphery is a reflection of changes in the brain, then in addition of potentially serving as a diagnostic tool, serum microRNAs may serve as a promising toolbox to decipher alcohol’s actions on the brain. Using bioinformatics tools, we identified several additional predicted targets of miR30a-5p, miR191-5p, miR195-5p and miR206-3p, all of which are expressed in the CNS (Table 1). For example, miR30a-5p is predicted to target GSKIP, a GSK3-beta and PKA scaffolding protein (Dema et al., 2016), and Rasd1, a member of the small G proteins, that plays a role in circadian rhythm (Cheng et al., 2006). miR191-5p is predicted to target actin and microtubules binding proteins including tropomodulin 2 which has been associated with synaptic plasticity and memory (Cox et al., 2003; Omotade et al., 2018). miR195-5p targets MAPK8, a member of the MAP kinase family (Davis, 2000), and miR206-3p predicted targets include the actin binding protein, Coro1c (Roadcap et al., 2008), and complexin 2, a SNARE machinery gene (Trimbuch and Rosenmund, 2016). Thus, it would be of interest to explore the contribution of these targets and others to the Rapid Onset and/or the development of heavy alcohol use.

**Table 1.**
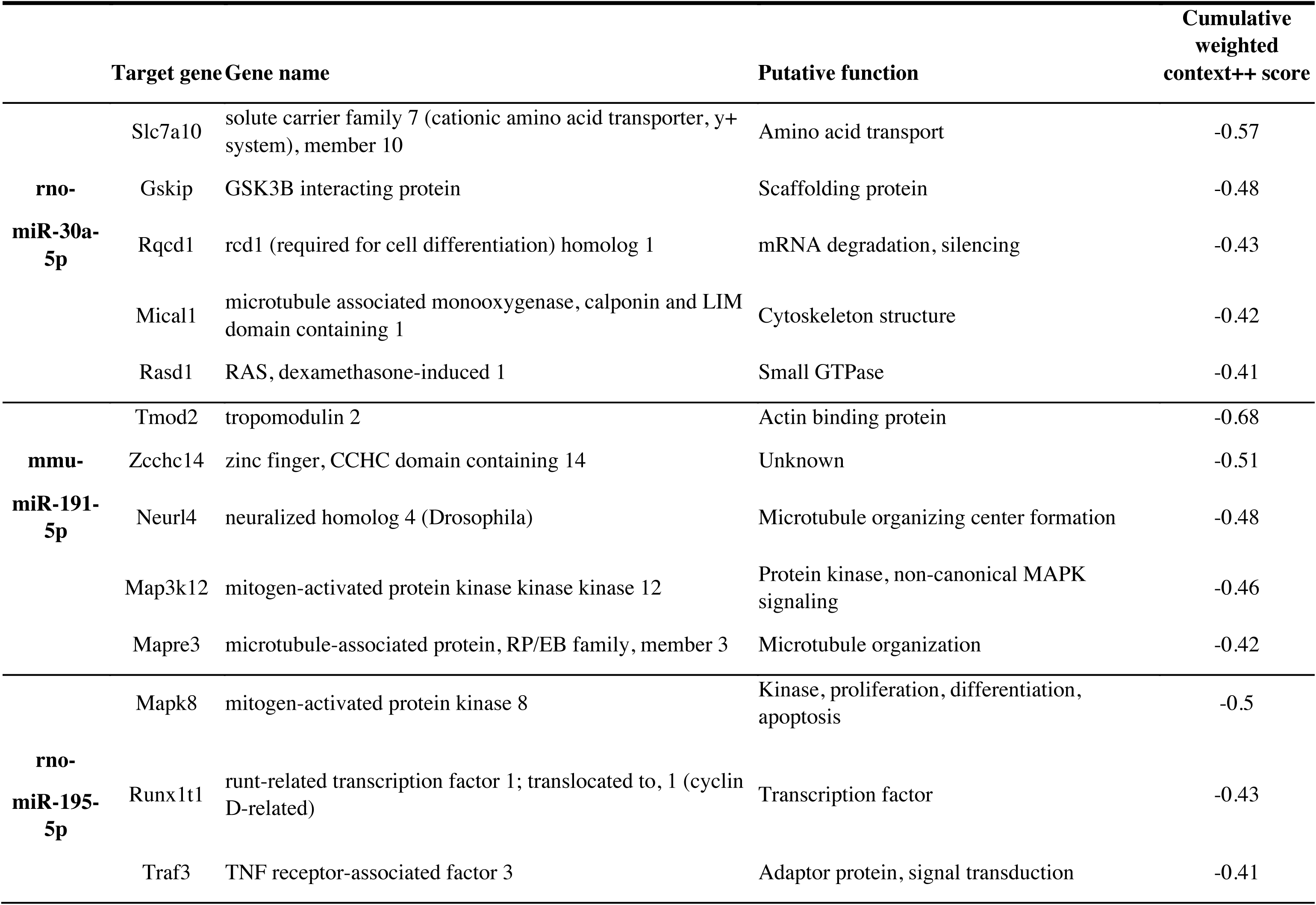
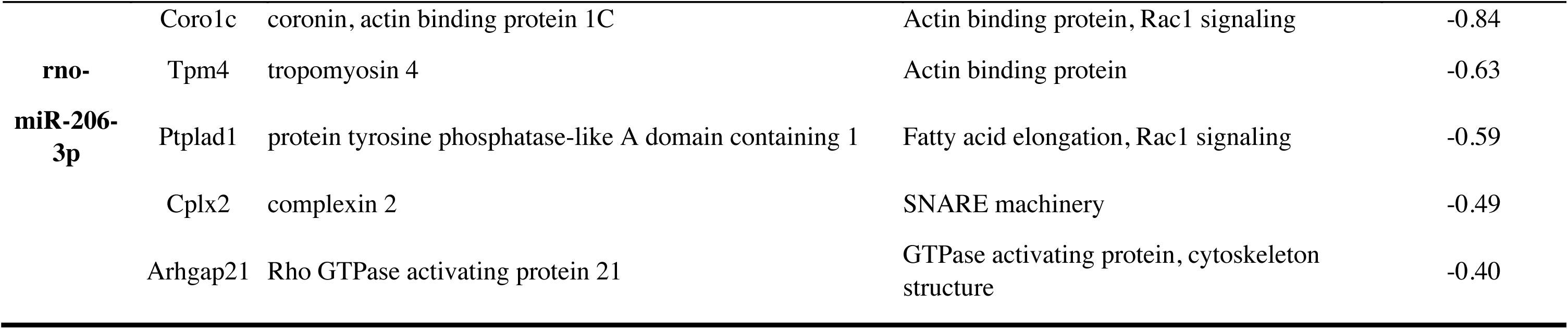
Predicted mRNA targets of miR-30a-5p, miR-191-5p, miR-195-5p and miR-206-3p. TargetScan database (http://www.targetscan.org/mmu_72/) was used to identify mRNAs predicted to be targeted by microRNA families containing each of the serum microRNA sequences. Rat (rno) sequences were used for miR195-5p, miR206-3p and miR30a-5p. miR191-5p mouse (mmu) sequence was used as the rat sequence was unavailable in TargetScan. Gene candidates predicted to be targeted by the specific microRNAs are ordered according to the cumulative weighted context++ score (−1 to 1). A lower score indicates higher predicted repression of the mRNA by the microRNA, considering 14 features of both the microRNA and the putative mRNA target, in addition to the seed site complementarity .

### Intermittent access to 20% alcohol does not alter platelet number

Numerous reports indicate that chronic alcohol use reduces platelet count in human alcoholics (Mikhailidis et al., 1986; Renaud and Ruf, 1996; Rubin and Rand, 1994). In contrast, we did not detect changes in platelet number in response to long-term alcohol use in rats. Taking into account that the BAC of rats drinking an average of 3-4 g/kg is in the range of 80 mg/dl, whereas the *in vitro* reduction in platelet number required concentrations of 200 mg/dl or higher (Mikhailidis et al., 1986; Rubin and Rand, 1994), it is not surprising that alcohol did not alter platelet number in our model. In addition, *in vitro* studies suggest an inhibitory effect of alcohol on platelet aggregation (Mikhailidis et al., 1986; Rubin and Rand, 1994). Whether or not cycles of binge and withdrawal alter platelet function, particularly platelet secretion mechanisms of relevance to BDNF, is an open question that will be addressed in future research.

### Implications

In summary, we found that the drinking profiles in Long-Evans rats that were subjected to 8 weeks of IA20%2BC are not homogeneous. Only 55% of rats are heavy drinkers and a subset of heavy users exhibits a very Rapid Onset of drinking. We show that *BDNF* protein levels were reduced in Late Onset and Rapid Onset heavy drinkers, however, microRNAs targeting BDNF were found to be increased only in Rapid Onset animals. Thus, our data raise the possibility that BDNF targeting microRNAs may be developed as biomarkers for early detection of AUD. microRNAs have been shown to be potential biomarkers for numerous diseases (Cortez and Calin, 2009) including depression (Yuan et al., 2018). Thus, human research is warranted to determine whether serum microRNAs can be used as predictor of AUD, as well as tools to monitor the disease. Furthermore, intravenous administration of an inhibitor of microRNA miR122 reduced plasma cholesterol and fatty liver disease in mice (Cortez and Calin, 2009), raising an interesting possibility that microRNA inhibitors could be developed as new therapy for the treatment of AUD.

## Acknowledgments

This research was supported by the National Institute of Alcohol Abuse and Alcoholism R37AA01684 (D.R.). The authors thank Dr. Sebastien Carnicella for his input.

---

**Supplementary Figure 1.**
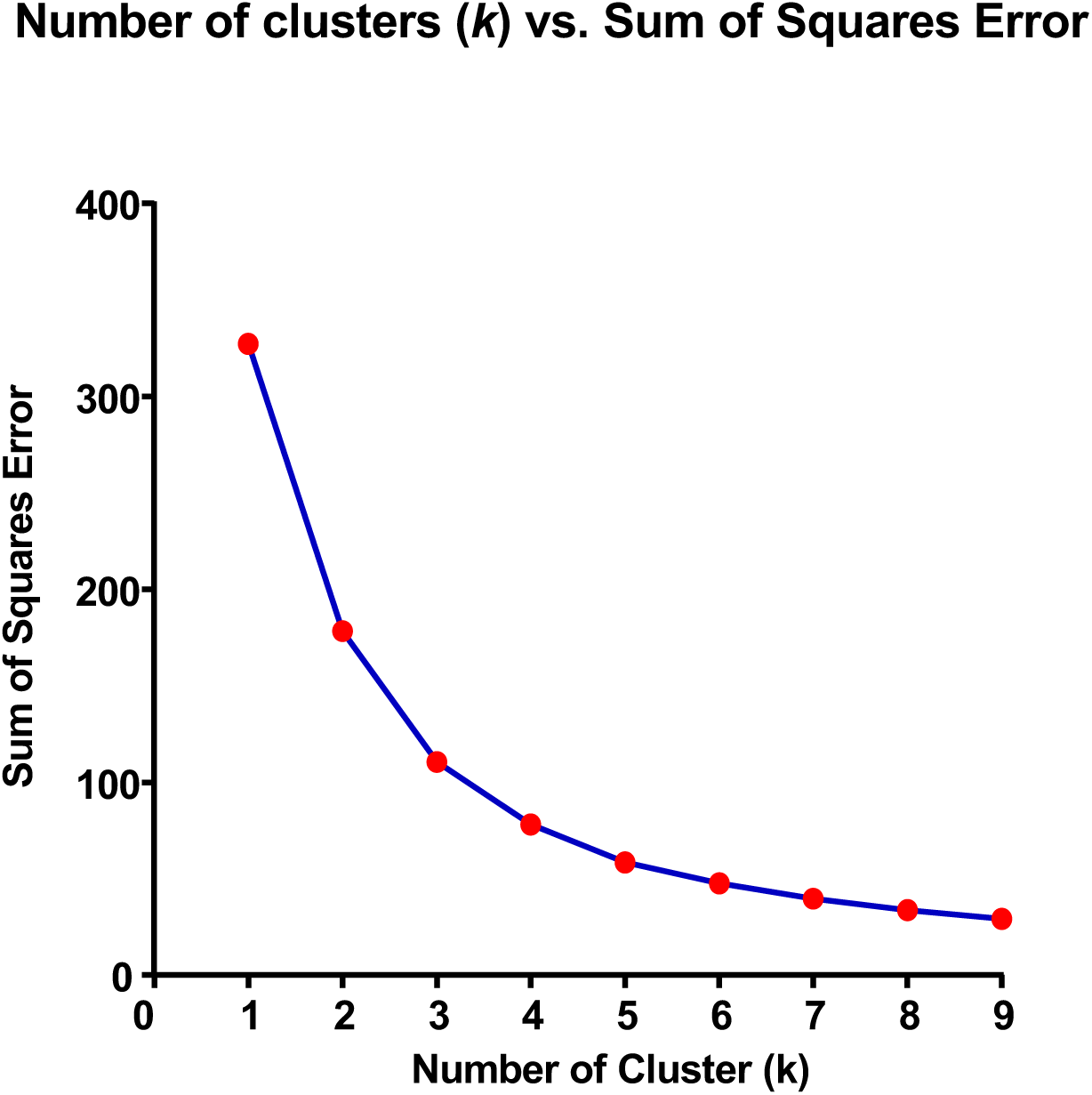
Number of clusters (k) vs SSE. Plot of number of clusters (*k*) vs. sum-of-squares error (SSE) for K-means clustering of alcohol intake (g/kg) averaged over the first four and last four drinking sessions (n=55). The decline in SSE plateauing at *k*=3 clusters supports the presence of three drinking phenotypes.

## References

Anitha A, Thanseem I (2015) microRNA and Autism. Adv Exp Med Biol 888:71–83.

Augier E, Barbier E, Dulman RS, Licheri V, Augier G, Domi E, Barchiesi R, Farris S, Natt D, Mayfield RD, Adermark L, Heilig M (2018) A molecular mechanism for choosing alcohol over an alternative reward. Science 360:1321–1326.

Bekinschtein P, Cammarota M, Medina JH (2014) BDNF and memory processing. Neuropharmacology 76 iPt C:677–683.

Bothwell M (2014) NGF, BDNF, NT3, and NT4. Handbook of experimental pharmacology 220:3–15.

Carnicella S, Amamoto R, Ron D (2009) Excessive alcohol consumption is blocked by glial cell line-derived neurotrophic factor. Alcohol 43:35–43.

Carnicella S, Ron D, Barak S (2014) Intermittent ethanol access schedule in rats as a preclinical model of alcohol abuse. Alcohol 48:243–252.

Castren E (2014) Neurotrophins and psychiatric disorders. Handb Exp Pharmacol 220:461–479.

Cattaneo A, Cattane N, Begni V, Pariante CM, Riva MA (2016) The human BDNF gene: peripheral gene expression and protein levels as biomarkers for psychiatric disorders. Transl Psychiatry 6:e958.

Ceballos N, Sharma S (2016) Risk and Resilience: The Role of Brain-derived Neurotrophic Factor in Alcohol Use Disorder. AIMS Neuroscience 3:398–432.

Chacon-Fernandez P, Sauberli K, Colzani M, Moreau T, Ghevaert C, Barde YA (2016) Brain-derived Neurotrophic Factor in Megakaryocytes. J Biol Chem 291:9872–9881.

Chandrasekar V, Dreyer JL (2009) microRNAs miR-124, let-7d and miR-181a regulate cocaine-induced plasticity. Mol Cell Neurosci 42:350–362.

Cheng HY, Dziema H, Papp J, Mathur DP, Koletar M, Ralph MR, Penninger JM, Obrietan K (2006) The molecular gatekeeper Dexras1 sculpts the photic responsiveness of the mammalian circadian clock. J Neurosci 26:12984–12995.

Cortez MA, Calin GA (2009) MicroRNA identification in plasma and serum: a new tool to diagnose and monitor diseases. Expert Opin Biol Ther 9:703–711.

Costa MA, Girard M, Dalmay F, Malauzat D (2011) Brain-derived neurotrophic factor serum levels in alcohol-dependent subjects 6 months after alcohol withdrawal. Alcoholism, clinical and experimental research 35:1966–1973.

Cox PR, Fowler V, Xu B, Sweatt JD, Paylor R, Zoghbi HY (2003) Mice lacking Tropomodulin-2 show enhanced long-term potentiation, hyperactivity, and deficits in learning and memory. Mol Cell Neurosci 23:1–12.

Darcq E, Morisot N, Phamluong K, Warnault V, Jeanblanc J, Longo FM, Massa SM, Ron D (2016) The Neurotrophic Factor Receptor p75 in the Rat Dorsolateral Striatum Drives Excessive Alcohol Drinking. J Neurosci 36:10116–10127.

Darcq E, Warnault V, Phamluong K, Besserer GM, Liu F, Ron D (2015) MicroRNA-30a-5p in the prefrontal cortex controls the transition from moderate to excessive alcohol consumption. Molecular psychiatry 20:1219–1231.

Davis RJ (2000) Signal transduction by the JNK group of MAP kinases. Cell 103:239–252.

Dema A, Schroter MF, Perets E, Skroblin P, Moutty MC, Deak VA, Birchmeier W, Klussmann E (2016) The A-Kinase Anchoring Protein (AKAP) Glycogen Synthase Kinase 3beta Interaction Protein (GSKIP) Regulates beta-Catenin through Its Interactions with Both Protein Kinase A (PKA) and GSK3beta. J Biol Chem 291:19618–19630.

Edwards AC, Gillespie NA, Aggen SH, Kendler KS (2013) Assessment of a modified DSM-5 diagnosis of alcohol use disorder in a genetically informative population. Alcoholism, clinical and experimental research 37:443–451.

Eichner ER (1973) The hematologic disorders of alcoholism. Am J Med 54:621–630.

Enoch MA, Goldman D (2002) Problem drinking and alcoholism: diagnosis and treatment. American family physician 65:441–448.

Fernandes BS, Gama CS, Cereser KM, Yatham LN, Fries GR, Colpo G, de Lucena D, Kunz M, Gomes FA, Kapczinski F (2011) Brain-derived neurotrophic factor as a state-marker of mood episodes in bipolar disorders: a systematic review and meta-regression analysis. J Psychiatr Res 45:995–1004.

Fujimura H, Altar CA, Chen R, Nakamura T, Nakahashi T, Kambayashi J, Sun B, Tandon NN (2002) Brain-derived neurotrophic factor is stored in human platelets and released by agonist stimulation. Thromb Haemost 87:728–734.

Garcia-Marchena N, Silva-Pena D, Martin-Velasco AI, Villanua MA, Araos P, Pedraz M, Maza-Quiroga R, Romero-Sanchiz P, Rubio G, Castilla-Ortega E, Suarez J, Rodriguez de Fonseca F, Serrano A, Pavon FJ (2017) Decreased plasma concentrations of BDNF and IGF-1 in abstinent patients with alcohol use disorders. PloS one 12:e0187634.

Gowin JL, Sloan ME, Stangl BL, Vatsalya V, Ramchandani VA (2017) Vulnerability for Alcohol Use Disorder and Rate of Alcohol Consumption. Am J Psychiatry 174:1094–1101.

Grant BF, Goldstein RB, Saha TD, Chou SP, Jung J, Zhang H, Pickering RP, Ruan WJ, Smith SM, Huang B, Hasin DS (2015) Epidemiology of DSM-5 Alcohol Use Disorder: Results From the National Epidemiologic Survey on Alcohol and Related Conditions III. JAMA psychiatry 72:757–766.

Heberlein A, Muschler M, Wilhelm J, Frieling H, Lenz B, Groschl M, Kornhuber J, Bleich S, Hillemacher T (2010) BDNF and GDNF serum levels in alcohol-dependent patients during withdrawal. Prog Neuropsychopharmacol Biol Psychiatry 34:1060–1064.

Huang EJ, Reichardt LF (2003) Trk receptors: roles in neuronal signal transduction. Annu Rev Biochem 72:609–642.

Ignacio C, Hicks SD, Burke P, Lewis L, Szombathyne-Meszaros Z, Middleton FA (2015) Alterations in serum microRNA in humans with alcohol use disorders impact cell proliferation and cell death pathways and predict structural and functional changes in brain. BMC Neurosci 16:55.

James G, Witten D, Hastie T, Tibshirani R (2013) Unsupervised Learning In: Introduction to Statistical Learning: with Applications in R. Springer-Verlag New York: New York N.Y.

Jeanblanc J, He DY, Carnicella S, Kharazia V, Janak PH, Ron D (2009) Endogenous BDNF in the dorsolateral striatum gates alcohol drinking. J Neurosci 29:13494–13502.

Jeanblanc J, He DY, McGough NN, Logrip ML, Phamluong K, Janak PH, Ron D (2006) The dopamine D3 receptor is part of a homeostatic pathway regulating ethanol consumption. J Neurosci 26:1457–1464.

Jeanblanc J, Logrip ML, Janak PH, Ron D (2013) BDNF-mediated regulation of ethanol consumption requires the activation of the MAP kinase pathway and protein synthesis. Eur J Neurosci 37:607–612.

Jin F, Hu H, Xu M, Zhan S, Wang Y, Zhang H, Chen X (2018) Serum microRNA Profiles Serve as Novel Biomarkers for Autoimmune Diseases. Front Immunol 9:2381.

Kermani P, Hempstead B (2019) BDNF Actions in the Cardiovascular System: Roles in Development, Adulthood and Response to Injury. Front Physiol 10:455.

Klimkiewicz A, Mach A, Jakubczyk A, Klimkiewicz J, Wnorowska A, Kopera M, Fudalej S, Burmeister M, Brower K, Wojnar M (2017) COMT and BDNF Gene Variants Help to Predict Alcohol Consumption in Alcohol-dependent Patients. J Addict Med 11:114–118.

Kraemer BR, Snow JP, Vollbrecht P, Pathak A, Valentine WM, Deutch AY, Carter BD (2014a) A role for the p75 neurotrophin receptor in axonal degeneration and apoptosis induced by oxidative stress. J Biol Chem 289:21205–21216.

Kraemer BR, Yoon SO, Carter BD (2014b) The biological functions and signaling mechanisms of the p75 neurotrophin receptor. Handb Exp Pharmacol 220:121–164.

Krahe TE, El-Danaf RN, Dilger EK, Henderson SC, Guido W (2011) Morphologically distinct classes of relay cells exhibit regional preferences in the dorsal lateral geniculate nucleus of the mouse. J Neurosci 31:17437–17448.

Leal G, Comprido D, Duarte CB (2014) BDNF-induced local protein synthesis and synaptic plasticity. Neuropharmacology 76 Pt C:639–656.

Linker RA, Lee DH, Demir S, Wiese S, Kruse N, Siglienti I, Gerhardt E, Neumann H, Sendtner M, Luhder F, Gold R (2010) Functional role of brain-derived neurotrophic factor in neuroprotective autoimmunity: therapeutic implications in a model of multiple sclerosis. Brain 133:2248–2263.

Linker RA, Lee DH, Flach AC, Litke T, van den Brandt J, Reichardt HM, Lingner T, Bommhardt U, Sendtner M, Gold R, Flugel A, Luhder F (2015) Thymocyte-derived BDNF influences T-cell maturation at the DN3/DN4 transition stage. Eur J Immunol 45:1326–1338.

Logrip ML, Barak S, Warnault V, Ron D (2015) Corticostriatal BDNF and alcohol addiction. Brain Res 1628:60–67.

Logrip ML, Janak PH, Ron D (2008) Dynorphin is a downstream effector of striatal BDNF regulation of ethanol intake. FASEB J 22:2393–2404.

Logrip ML, Janak PH, Ron D (2009) Escalating ethanol intake is associated with altered corticostriatal BDNF expression. J Neurochem 109:1459–1468.

Maffioletti E, Zanardini R, Gennarelli M, Bocchio-Chiavetto L (2014) Influence of clotting duration on brain-derived neurotrophic factor (BDNF) dosage in serum. Biotechniques 57:111–114.

McBride WJ, Rodd ZA, Bell RL, Lumeng L, Li TK (2014) The alcohol-preferring (P) and high-alcohol-drinking (HAD) rats--animal models of alcoholism. Alcohol 48:209–215.

McGough NN, He DY, Logrip ML, Jeanblanc J, Phamluong K, Luong K, Kharazia V, Janak PH, Ron D (2004) RACK1 and brain-derived neurotrophic factor: a homeostatic pathway that regulates alcohol addiction. J Neurosci 24:10542–10552.

Mellios N, Huang HS, Baker SP, Galdzicka M, Ginns E, Akbarian S (2009) Molecular determinants of dysregulated GABAergic gene expression in the prefrontal cortex of subjects with schizophrenia. Biological psychiatry 65:1006–1014.

Mellios N, Huang HS, Grigorenko A, Rogaev E, Akbarian S (2008) A set of differentially expressed miRNAs, including miR-30a-5p, act as post-transcriptional inhibitors of BDNF in prefrontal cortex. Hum Mol Genet 17:3030–3042.

Mikhailidis DP, Jenkins WJ, Barradas MA, Jeremy JY, Dandona P (1986) Platelet function defects in chronic alcoholism. Br Med J (Clin Res Ed) 293:715–718.

Minichiello L (2009) TrkB signalling pathways in LTP and learning. Nature reviews Neuroscience 10:850–860.

Mitchell PS, Parkin RK, Kroh EM, Fritz BR, Wyman SK, Pogosova-Agadjanyan EL, Peterson A, Noteboom J, O’Briant KC, Allen A, Lin DW, Urban N, Drescher CW, Knudsen BS, Stirewalt DL, Gentleman R, Vessella RL, Nelson PS, Martin DB, Tewari M (2008) Circulating microRNAs as stable blood-based markers for cancer detection. Proceedings of the National Academy of Sciences of the United States of America 105:10513–10518.

Molendijk ML, Bus BA, Spinhoven P, Penninx BW, Kenis G, Prickaerts J, Voshaar RC, Elzinga BM (2011) Serum levels of brain-derived neurotrophic factor in major depressive disorder: state-trait issues, clinical features and pharmacological treatment. Mol Psychiatry 16:1088–1095.

Nakahashi T, Fujimura H, Altar CA, Li J, Kambayashi J, Tandon NN, Sun B (2000) Vascular endothelial cells synthesize and secrete brain-derived neurotrophic factor. FEBS Lett 470:113–117.

Nees F, Witt SH, Dinu-Biringer R, Lourdusamy A, Tzschoppe J, Vollstadt-Klein S, Millenet S, Bach C, Poustka L, Banaschewski T, Barker GJ, Bokde AL, Bromberg U, Buchel C, Conrod PJ, Frank J, Frouin V, Gallinat J, Garavan H, Gowland P, Heinz A, Ittermann B, Mann K, Martinot JL, Paus T, Pausova Z, Robbins TW, Smolka MN, Rietschel M, Schumann G, Flor H, consortium I (2015) BDNF Val66Met and reward-related brain function in adolescents: role for early alcohol consumption. Alcohol 49:103–110.

Nubukpo P, Ramoz N, Girard M, Malauzat D, Gorwood P (2017) Determinants of Blood Brain-Derived Neurotrophic Factor Blood Levels in Patients with Alcohol Use Disorder. Alcoholism, clinical and experimental research 41:1280–1287.

Omotade OF, Rui Y, Lei W, Yu K, Hartzell HC, Fowler VM, Zheng JQ (2018) Tropomodulin Isoform-Specific Regulation of Dendrite Development and Synapse Formation. J Neurosci 38:10271–10285.

Pandey SC, Zhang H, Roy A, Misra K (2006) Central and medial amygdaloid brain-derived neurotrophic factor signaling plays a critical role in alcohol-drinking and anxiety-like behaviors. J Neurosci 26:8320–8331.

Panja D, Bramham CR (2014) BDNF mechanisms in late LTP formation: A synthesis and breakdown. Neuropharmacology 76 Pt C:664–676.

Park H, Poo MM (2013) Neurotrophin regulation of neural circuit development and function. Nature reviews Neuroscience 14:7–23.

Polyakova M, Stuke K, Schuemberg K, Mueller K, Schoenknecht P, Schroeter ML (2015) BDNF as a biomarker for successful treatment of mood disorders: a systematic & quantitative meta-analysis. Journal of affective disorders 174:432–440.

Prakash A, Zhang H, Pandey SC (2008) Innate differences in the expression of brain-derived neurotrophic factor in the regions within the extended amygdala between alcohol preferring and nonpreferring rats. Alcohol Clin Exp Res 32:909–920.

Qureshi IA, Mehler MF (2012) Emerging roles of non-coding RNAs in brain evolution, development, plasticity and disease. Nature reviews Neuroscience 13:528–541.

Raivio N, Miettinen P, Kiianmaa K (2014) Innate BDNF expression is associated with ethanol intake in alcohol-preferring AA and alcohol-avoiding ANA rats. Brain research 1579:74–83.

Renaud SC, Ruf JC (1996) Effects of alcohol on platelet functions. Clin Chim Acta 246:77–89.

Rizos EN, Rontos I, Laskos E, Arsenis G, Michalopoulou PG, Vasilopoulos D, Gournellis R, Lykouras L (2008) Investigation of serum BDNF levels in drug-naive patients with schizophrenia. Prog Neuropsychopharmacol Biol Psychiatry 32:1308–1311.

Roadcap DW, Clemen CS, Bear JE (2008) The role of mammalian coronins in development and disease. Subcell Biochem 48:124–135.

Ron D, Berger A (2018) Targeting the intracellular signaling “STOP” and “GO” pathways for the treatment of alcohol use disorders. Psychopharmacology (Berl) 235:1727–1743.

Rubin R, Rand ML (1994) Alcohol and platelet function. Alcoholism, clinical and experimental research 18:105–110.

Ruiz CR, Shi J, Meffert MK (2014) Transcript specificity in BDNF-regulated protein synthesis. Neuropharmacology 76 Pt C:657–663.

Silva-Pena D, Garcia-Marchena N, Alen F, Araos P, Rivera P, Vargas A, Garcia-Fernandez MI, Martin-Velasco AI, Villanua MA, Castilla-Ortega E, Santin L, Pavon FJ, Serrano A, Rubio G, Rodriguez de Fonseca F, Suarez J (2019) Alcohol-induced cognitive deficits are associated with decreased circulating levels of the neurotrophin BDNF in humans and rats. Addiction biology 24:1019–1033.

Sunderland N, Skroblin P, Barwari T, Huntley RP, Lu R, Joshi A, Lovering RC, Mayr M (2017) MicroRNA Biomarkers and Platelet Reactivity: The Clot Thickens. Circ Res 120:418–435.

Tapocik JD, Barbier E, Flanigan M, Solomon M, Pincus A, Pilling A, Sun H, Schank JR, King C, Heilig M (2014) microRNA-206 in rat medial prefrontal cortex regulates BDNF expression and alcohol drinking. J Neurosci 34:4581–4588.

Teng HK, Teng KK, Lee R, Wright S, Tevar S, Almeida RD, Kermani P, Torkin R, Chen ZY, Lee FS, Kraemer RT, Nykjaer A, Hempstead BL (2005) ProBDNF induces neuronal apoptosis via activation of a receptor complex of p75NTR and sortilin. J Neurosci 25:5455–5463.

Torres-Berrio A, Nouel D, Cuesta S, Parise EM, Restrepo-Lozano JM, Larochelle P, Nestler EJ, Flores C (2019) MiR-218: a molecular switch and potential biomarker of susceptibility to stress. Mol Psychiatry. doi: 10.1038/s41380-019-0421-5. [Epub ahead of print].

Trimbuch T, Rosenmund C (2016) Should I stop or should I go? The role of complexin in neurotransmitter release. Nature reviews Neuroscience 17:118–125.

Warnault V, Darcq E, Morisot N, Phamluong K, Wilbrecht L, Massa SM, Longo FM, Ron D (2016) The BDNF Valine 68 to Methionine Polymorphism Increases Compulsive Alcohol Drinking in Mice That Is Reversed by Tropomyosin Receptor Kinase B Activation. Biological psychiatry 79:463–473.

WHO (2014) World Health Statistics 2014. World Health Organization. pp 1-180.

Wojnar M, Brower KJ, Strobbe S, Ilgen M, Matsumoto H, Nowosad I, Sliwerska E, Burmeister M (2009) Association between Val66Met brain-derived neurotrophic factor (BDNF) gene polymorphism and post-treatment relapse in alcohol dependence. Alcohol Clin Exp Res 33:693–702.

Woo NH, Teng HK, Siao CJ, Chiaruttini C, Pang PT, Milner TA, Hempstead BL, Lu B (2005) Activation of p75NTR by proBDNF facilitates hippocampal long-term depression. Nature neuroscience 8:1069–1077.

Yuan H, Mischoulon D, Fava M, Otto MW (2018) Circulating microRNAs as biomarkers for depression: Many candidates, few finalists. J Affect Disord 233:68–78.

Zagrebelsky M, Holz A, Dechant G, Barde YA, Bonhoeffer T, Korte M (2005) The p75 neurotrophin receptor negatively modulates dendrite complexity and spine density in hippocampal neurons. J Neurosci 25:9989–9999.

Zanardini R, Fontana A, Pagano R, Mazzaro E, Bergamasco F, Romagnosi G, Gennarelli M, Bocchio-Chiavetto L (2011) Alterations of brain-derived neurotrophic factor serum levels in patients with alcohol dependence. Alcoholism, clinical and experimental research 35:1529–1533.

Zhou L, Xiong J, Ruan CS, Ruan Y, Liu D, Bao JJ, Zhou XF (2018) ProBDNF/p75NTR/sortilin pathway is activated in peripheral blood of patients with alcohol dependence. Transl Psychiatry 7:2.

